# Surgically disconnected temporal pole exhibits resting functional connectivity with remote brain regions

**DOI:** 10.1101/127571

**Authors:** David E. Warren, Matthew J. Sutterer, Joel Bruss, Taylor J. Abel, Andrew Jones, Hiroto Kawasaki, Michelle Voss, Martin Cassell, Matthew A. Howard, Daniel Tranel

## Abstract

Functional connectivity, as measured by resting-state fMRI, has proven a powerful method for studying brain systems in the context of behavior, development, and disease states. However, the relationship of functional connectivity to structural connectivity remains unclear. If functional connectivity relies on structural connectivity, then anatomical isolation of a brain region should eliminate functional connectivity with other brain regions. We tested this by measuring functional connectivity of the surgically disconnected temporal pole in resection patients (N=5; mean age 37; 2F, 3M). Functional connectivity was evaluated based on coactivation of whole-brain fMRI data with the average low-frequency BOLD signal from disconnected tissue in each patient. In sharp contrast to our prediction, we observed significant functional connectivity between the disconnected temporal pole and remote brain regions in each disconnection case. These findings raise important questions about the neural bases of functional connectivity measures derived from the fMRI BOLD signal.

## 1. Introduction

THe current consensus understanding of functional connectivity measured with resting-state functional magnetic resonance imaging (fMRI) is that measures of functional connectivity reflect — in part — temporal coactivation between neuronal populations. This follows from two premises. First, temporally correlated fMRI signals between brain regions reflect coordinated neural activity. Converging evidence from intracranial recordings, EEG, and fMRI supports this premise (David et al., 2008; Logothetis, Pauls, Augath, Trinath, & Oeltermann, 2001). Second, a necessary condition of functional connectivity that reflects communication between neuronal populations is structural connectivity, whether monosynaptic or polysynaptic (e.g., Buckner, Krienen, & Yeo, 2013). This premise yields the straightforward prediction that a brain region with no structural connections to other brain regions would be functionally isolated. Some empirical support for this prediction exists (Gillebert & Mantini, 2013; Gre-fkes & Fink, 2011; Lu et al., 2011; Putnam, Wig, Grafton, Kelley, & Gazzaniga, 2008; Shen et al., 2015; Tovar-Moll et al., 2014; Tyszka, Kennedy, Adolphs, & Paul, 2011), but the strongest test of this prediction has not been conducted. Specifically, can brain tissue that has been fully, surgically disconnected from the rest of the brain show functional connectivity with other brain regions?

Prior efforts that addressed the complex relationship between structural and functional connectivity have provided some insight, but the implications of their findings are sometimes contradictory (Damoiseaux & Greicius, 2009; Greicius, Supekar, Menon, & Dougherty, 2009; Shen et al., 2015). Structural connectivity does predict some aspects of functional connectivity (Adachi et al., 2012; Honey et al., 2009), and changes in structural connectivity have been shown to alter functional connectivity (Gillebert & Mantini, 2013; Grefkes & Fink, 2011; Lu et al., 2011; Putnam et al., 2008; Shen et al., 2015; Tovar-Moll et al., 2014; Tyszka et al., 2011). In one example of function changing with structure, adult macaques with corpus callosotomy showed radical changes in interhemispheric functional connectivity (O’Reilly et al., 2013). Similarly, pontine strokes have been shown to influence functional connectivity of damaged cerebellar-cortical circuits (Lu et al., 2011). Findings of this nature would be consistent with a simple model of structural-to-functional connectivity, but other findings present challenges for such a model. In a healthy brain, very strong functional connectivity of bilateral primary visual cortex occurs in the absence of monosynaptic structural connections (Vincent et al., 2007). Even the interruption of major structural connections such as the corpus callosum does not necessarily abolish functional connectivity as reported from human studies of individuals with developmental callosal agenesis (Tyszka et al., 2011) or receiving corpus callosotomy (Uddin et al., 2008; but see Johnston et al., 2008).

The diversity of findings relating structural to functional connectivity reinforces the importance of rigorously evaluating the basic premise that structural connectivity is necessary for functional connectivity that reflects communication between neuronal populations. An important study of neuroimaging data from healthy adults characterized the relationship between functional and structural connectivity but relied on diffusion-weighted neuroimaging data to draw inferences regarding structural connectivity (Honey et al., 2009). Alternatively, in neurological cases with altered or abolished structural connectivity of the corpus callosum, the observed functional connectivity may be supported by other structural connections such as ascending innervation, indirect structural connectivity within the cerebral cortex, or both (Tyszka et al., 2011; Uddin et al., 2008). Studies of individuals with pontine stroke (Lu et al., 2011) have relied on organically-occurring brain lesions that were not subject to any artificial control or direct in vivo inspection. Addressing whether intentional structural disconnection of human brain tissue leads to functional isolation would provide fundamental insight into the mechanisms driving functional connectivity as reflected from the fMRI BOLD signal with implications for variance not accounted for by structural connectivity (Honey et al., 2009) or known sources of noise such as motion and physiology (Power, Barnes, Snyder, Schlaggar, & Petersen, 2012; Satterthwaite et al., 2012; Shmueli et al., 2007; Koene R. A. Van Dijk, Sabuncu, & Buckner, 2012; Weisskoff et al., 1993).

In the current study, we used resting-state fMRI to study the functional connectivity of a brain region that was anatomically isolated from the rest of the brain by surgical resection. We recruited patients who underwent unilateral temporal disconnection (TD) surgery to treat pharmacoresistant temporal lobe epilepsy (N=5; Table 1). In this surgical procedure, a portion of the temporal pole — including its pial vascular supply — is preserved but structurally isolated from the rest of the brain. Patients with TD provide a compelling opportunity to test the key premise that structural connectivity is necessary for functional connectivity. More specifically, we tested the prediction that surgical isolation of the temporal pole would result in no functional connectivity with any brain regions. We addressed this prediction by evaluating fMRI data from patients with TD using several approaches, including seed-based analysis, data-driven independent component analysis, and comparisons of spatial similarity of functional connectivity with normative expectations. Null results for evidence of functional connectivity would be consistent with our prediction that structural disconnection prevented functional connectivity. Alternatively, evidence of functional connectivity between the disconnected temporal pole and other brain regions would instead be consistent with the proposition that structural connectivity is not necessary for functional connectivity in fMRI BOLD signal.

## 2. Methods

### 2.1 Participants

#### 2.1.1 Selection procedure

We recruited 6 participants (2F/4M) from the Iowa Neurological Patient Registry who previously underwent unilateral TD (four left, two right) for the treatment of pharmacoresistant epilepsy. The research was approved by the Human Subjects Committee of the University of Iowa Institutional Review Board, and all participants gave informed consent to participate in research. These participants were identified by searching the Registry: the initial search found 57 active Registry cases who underwent surgery for temporal lobe epilepsy who also had appropriate MRI imaging. This pool of 57 cases was then screened for temporal pole disconnections using anatomical criteria described below. Two neurosurgeons (HK & TA) inspected the anatomical T1 imaging for each case and had to reach agreement that the case was a true disconnection for inclusion in this project. This screening excluded 45 cases, leaving 12 cases with confirmed disconnection. All 12 cases met anatomical criteria based on microscopic inspection during surgery and evaluation of structural MRI data collected in the chronic epoch. These 12 cases were further screened for those who had > 15 minutes of resting-state fMRI data because that duration has been associated with stabilization of intra-individual resting-state functional connectivity patterns (Laumann et al., 2016). Six cases were excluded for having insufficient resting state data, and this yielded the reported sample of 6 participants. One of these participants (3712L) was later excluded to due excessive motion during MRI exam (see below), yielding the final sample of participants (N=5).

**Table 1:** Demographic and neuropsychological data for temporal disconnection (TD) participants.

| ID | Sex | Hand | Age | Edu | Epoch | FSIQ | VCI | PRI | WRAT | Trails | AVLT | CFT | BNT | COWA |
| --- | --- | --- | --- | --- | --- | --- | --- | --- | --- | --- | --- | --- | --- | --- |
| 3360L | F | 100 | 29 | 16 | pre | 98<sup>a</sup> | — | — | — | 21/44 | 6/15/12 | 27/7.5 | 55 | 58 |
|  |  |  |  |  | post | 104<sup>b</sup> | 98 | 109 | 94 | 14/58 | 6/11/7 | 34/24<sup>c</sup> | 48 | 52 |
| 3712L* | M | 100 | 32 | 12 | pre | 82 | 87 | 88 | 74 | 48/90 | 6/9/4 | 33/16.5 | 36 | 23 |
|  |  |  |  |  | post | 89 | 93 | 100 | 76 | 50/99 | 5/8/3 | 35/10 | 43 | 30 |
| 3749L | M | -90 | 41 | 18 | pre | 92 | 100 | 90 | 85 | 16/49 | 7/7/4 | 29/15 | 45 | 35 |
|  |  |  |  |  | post | — | — | — | — | — | — | — | — | — |
| 3786L | F | -100 | 49 | 14 | pre | 84 | 96 | 75 | 88 | 36/161 | 5/13/8 | 27/11 | 56 | 45 |
|  |  |  |  |  | post | 81 | 96 | 79 | 87 | 40/129 | 3/7/5 | 26/9 | 57 | 44 |
| 3753R | M | 100 | 20 | 14 | pre | 105 | 112 | 111 | 112 | 33/59 | 5/12/12 | 30/21 | 53 | 32 |
|  |  |  |  |  | post | — | — | — | — | — | — | — | — | — |
| 3794R | M | 100 | 32 | 16 | pre | 104 | 87 | 117 | 96 | 16/33 | 6/12/8 | 29.5/15.5 | 46 | 26 |
|  |  |  |  |  | post | 104 | 91 | 105 | 96 | 15/29 | 8/10/10 | 31.5/27<sup>c</sup> | 50 | 26 |
| mean |  |  | 33.8 | 15.0 | pre | 94.2 | 96.4 | 96.2 | 91.0 | 28.3/72.7 | 5.8/11.3/8.0 | 29.2/14.4 | 48.5 | 36.5 |
| s.d. |  |  | 10.0 | 2.1 |  | 9.8 | 10.4 | 17.4 | 14.1 | 12.8/47.4 | 0.8/2.9/3.6 | 2.2/4.7 | 7.7 | 13.0 |
| mean |  |  |  |  | post | 94.5 | 94.5 | 98.2 | 88.2 | 29.8/78.8 | 5.5/9.0/6.2 | 31.6/17.5 | 49.5 | 38.0 |
| s.d. |  |  |  |  |  | 11.4 | 3.1 | 13.4 | 9.0 | 18.1/44.1 | 2.1/1.8/3.0 | 4.0/9.3 | 5.8 | 12.1 |

Note: Top row, presurgical evaluation; bottom row, postsurgical evaluation. Mean and standard deviations summarize available scores from each epoch. Note that Abbreviations: Age, years; Hand, handedness (+100=fully right handed, -100=fully left handed); Edu., education, years; FSIQ, WAIS-IV full-scale IQ except where noted; VCI, verbal comprehension index (from WAIS-IV); PRI, perceptual reasoning index (from WAIS-IV); WRAT, Wide Range Achievement Test, Reading test performance (normative percentile); Trails, Trails A/B, seconds; AVLT, Rey Auditory Verbal Learning Test, trial 1/5/30-min. delayed recall; CFT, Complex Figure Test copy/recall; BNT, Boston Naming Test; COWA, Controlled Oral Word Association test. Symbols: \*, 3712L was excluded later for motion (see Methods); <sup>a</sup>, WAIS-III; <sup>b</sup>, IQ was measured at chronicity of 10 years; <sup>c</sup>, Taylor complex figure.

#### 2.1.2 Temporal lobe disconnection procedure and anatomy

TD participants had partial resections including portions of the anterior temporal lobe neocortex and white matter resulting in complete anatomical disconnection of residual anterior temporal lobe tissue. Regarding the procedure, the TD approach represents a modification of the anterior temporal lobectomy operation and has the advantages of a shortened procedure time, decreased intraoperative blood loss, and spares the normal venous vasculature of the temporal pole region. Medically, the goal of the TD procedure is to reduce the morbidity of temporal lobectomy by decreasing the overall amount of cortex that is resected in order to limit potential complications. Additionally, the disconnection procedure preserves cortical venous drainage, and this may also reduce the potential for postoperative complications.

### 2.2 Data collection

All data for this study were collected in the chronic epoch at least 6 months after a participant’s surgery.

#### 2.2.1 Neuropsychological data

All participants completed a battery of neuropsychological tests as part of their enrollment in the Patient Registry. Table 1 summarizes demographic and neuropsychological data for the TD participants.

#### 2.2.2 Neuroimaging data

All images were collected on a 3T head-only Siemens TIM Trio MRI scanner with a receive-only 12-channel phased-array head-coil. These imaging data were acquired as part of ongoing studies in the Departments of Neurosurgery and Neurology at the University of Iowa. As such, the anatomical sequence and the number of resting-state volumes collected differed slightly between participants. The number of resting-state volumes collected for each participant are indicated in Table 2. Structural imaging for participants 3360, 3712, 3786 consisted of one three-dimensional 1 mm isotropic coronal T1 MPRAGE (Magnetization Prepared Rapid Gradient Echo Imaging) protocol with repetition time (TR)=2530 ms, TE=3.04 ms, TI=900 ms, flip angle=10°, FOV=256×240×256 mm). A parallel imaging technique known as generalized autocalibrating partially parallel acquisition (GRAPPA) with an acceleration factor of two was used for the MPRAGE scan. Structural imaging for participants 3749, 3753, and 3794 consisted of one threedimensional 1 mm isotropic multi-echo MPRAGE protocol with TR=2530 ms, TE=(1.74 + 1.86×n ms, n={0,1,2,3}, TI=126 ms, flip angle=10°, Bandwidth=651 Hz/pixel, FOV=176×256×256 mm, GRAPPA acceleration factor of two.

**Table 2:**
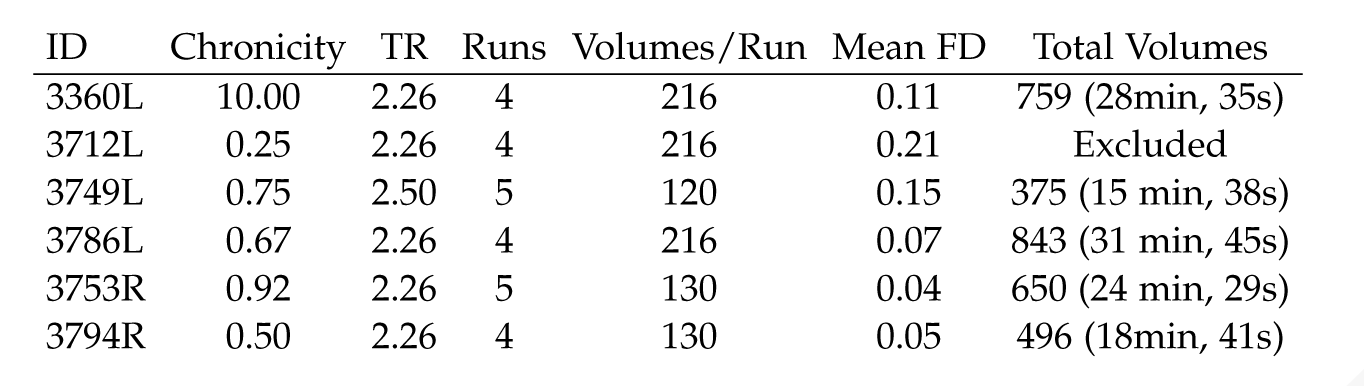
Functional neuroimaging parameters and quantities.

Functional neuroimaging of the resting blood oxygen level dependent (BOLD) effect was acquired with single-shot echo planar imaging (EPI). For all participants except 3749, multiple runs of resting-state fMRI data were obtained from each participant with whole brain coverage and the following scan parameters: slice dimensions=3.4375×3.4375×4.000 mm, TR=2260 ms, TE=30 ms, flip angle=80°, dwell=0.56, FOV=220×220×144 mm. For participant 3749, multiple runs of resting-state fMRI data were collected with whole brain coverage and the following scan parameters: slice dimensions=3.4375×3.4375×5.0000 mm, TR=2500 ms TE=30 ms, flip angle=80°, dwell=0.56, FOV=220×220×195 mm.

In addition to data collected from the TD participants at the University of Iowa, we drew on a publicly available dataset provided in the Cambridge Buckner Release of the 1000 Functional Connectomes Project (http://fcon_1000.projects.nitrc.org/) (Biswal et al., 2010). For comparison to our disconnected participants, we selected data from 25 participants who displayed minimal motion (see criteria below). The scanning parameters for these data were broadly similar to those used in the University of Iowa dataset and were as follows: slice dimensions=3×3×3 mm, TR=3000 ms, 119 volumes, TE=35 ms, dwell=0.70, FOV=216×216×141 mm.

### 2.3 Data analysis

#### 2.3.1 Structural MRI

T1 structural MRI data were used to evaluate the disconnection (TD) of temporal cortex (see above) and to manually generate masks for the lesion and the disconnected tissue.

Masks of the lesion cavity and disconnected tissue were manually generated from the native T1 anatomical images by authors AJ, MS, JB, and DW using FS-Lview (Jenkinson, Beckmann, Behrens, Woolrich, & Smith, 2012). All personnel involved in tracing were aware of the study’s broad goals, but mask inclusion or exclusion was governed solely by neuroanatomy and imaging data. All masks were revised until consensus was achieved among all personnel involved in mask generation. Masks of lesion and temporal pole were spatially adjacent but did not overlap. Masks of the temporal pole included all remaining gray and white matter of the disconnected region, and masks of the lesion included the space between the disconnected region and the preserved portion of the anterior temporal lobe. Temporal pole masks were used as a region-of-interest (ROI) for seed-based functional connectivity analyses, while lesion masks were used as a covariate seed to control for potentially confounding signal from the lesion cavity. For seeding in normative data, and for visualization of group overlap, native-space masks in T1 space were warped into standard space (ICBM MNI152) using FSL software (v5.0.2.2) nonlinear registration package FNIRT (FMRIB’s Software Library) (Andersson, Jenkinson, Smith, & others, 2007; Jenkinson et al., 2012). Following transformation to standard space, each mask was visually inspected and edited as necessary to ensure anatomical accuracy of the transformation. Structural MRI and disconnection masks of the TD participants are shown in Figure 1.

**Figure 1:**
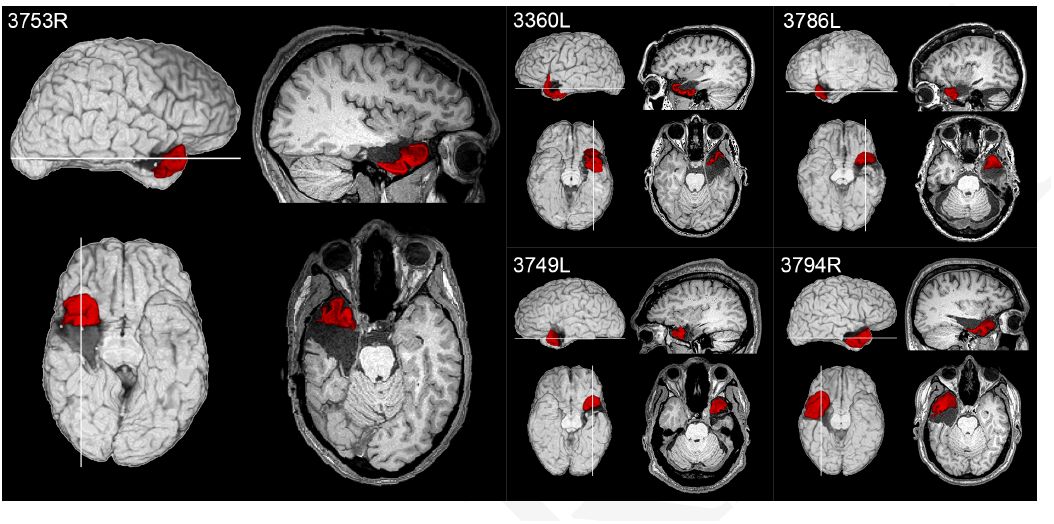
Anatomical images of temporal pole disconnection participants. Area highlighted in red represents the disconnection mask for each participant. L suffix for participant ID indicates left-sided disconnection, R indicates right-sided disconnection. Brain images are in radiological convention (i.e. left hemisphere is shown on the right side of the image).

#### 2.3.2 Functional MRI

fMRI data were processed using an approach typical of previous published work. Specifically, analyses were conducted in FSL (FMRIB’s Software Library 5.0.2.2), AFNI (Cox, 1996), and MATLAB (2015a, The MathWorks, Inc.) using established preprocessing and analytical steps for resting state data (Rigon, Duff, McAuley, Kramer, & Voss, 2016; K. R. A. Van Dijk et al., 2010; Voss et al., 2010, 2016). Each participant’s scan data underwent brain extraction, motion correction (3DVOLREG command from AFNI), and spatial smoothing (6 mm full-width-half-maximum kernel). Temporal filtering (3DBANDPASS function from AFNI) was done to ensure the BOLD fluctuations were within the frequency band of 0.008 < f < 0.08 Hz that has been identified as optimally sensitive to functional connectivity associated with brain networks (K. R. A. Van Dijk et al., 2010). Following preprocessing, further potential sources of noise were corrected for by extracting the mean time series from white matter, cerebral spinal fluid, and a global brain mask (representing the mean whole brain signal) (Fox, Zhang, Snyder, & Raichle, 2009; Power et al., 2014). In addition, the six head motion parameters described above were bandpassed with the same temporal filter applied to the fMRI data and included as nuisance regressors (Hallquist, Hwang, & Luna, 2013). Together, the 9 bandpassed nuisance regressors (white matter, CSF, global, and 6 motion parameters) were entered into a multiple regression as independent variables predicting the preprocessed rs-fMRI data (using FSL’s FEAT tool). Finally, because our region-of-interest was immediately adjacent to a lesion cavity, we also regressed the bandpassed mean time series from the lesion mask in each TD participant to eliminate local signals of no interest. This nuisance regressor was unique to the TD group.

Because motion is a particular concern in resting-state fMRI analyses we employed an additional stringent motion-censoring threshold recommended in the literature (Power et al., 2014). Specifically, we calculated the framewise displacement (FD) for each time-point in each resting-state run based on the motion-correction alignment, and we censored (i.e., deleted) frames where FD > 0.2 mm. To avoid instability in RSFC analyses, we excluded runs in which less than 50% of the data remained after motion censoring was applied. This criterion resulted in the complete exclusion of one participant, 3712, who had excessive motion in each of the runs of their fMRI data, and led to the exclusion of one of five runs for participant 3749. The final numbers of fMRI volumes after motion scrubbing for each participant are reported in Table 2. The residual fMRI data after nuisance regression and motion censoring was then used for the analyses described in the next section.

##### 2.3.2.1 ROI seed analysis

For each run of resting-state in each TD participant, the native-space disconnection mask for each TD participant, respectively, was used as a seed ROI. A statistical map was created by cross-correlating the seed time series with every other voxel in the brain (computed in MATLAB). These maps of Pearson’s correlations were transformed using Fisher’s r-to-Z transform to impose a normal distribution. Seed maps from individual resting-state runs were forwarded to a fixed effects analysis. A one-way t-test was calculated in AFNI’s 3DTTEST++ (as implemented in AFNI version 16.2.18) comparing the average of the individual run seed correlation map with zero. The t-statistic maps were transformed to Z-scores, and multiple comparisons for the resulting statistical maps were controlled using FSL’s CLUSTER command by thresholding the participant’s average seed map at Z > 2.33, with cluster correction of p < .05 (Worsley, Evans, Marrett, & Neelin, 1992). While we initially chose a lower Z-threshold to minimize the chance of false negative findings, we also examined the results of this analysis with a more stringent threshold of Z > 3.1 and the same cluster correction of p < .05. This procedure generated statistically thresholded maps for each participant showing the voxels that were reliably functionally connected with the disconnection seed ROI across the multiple resting-state runs. Finally, for comparison to each disconnection case, the standard MNI space disconnection mask for that TD participant was used in a similar ROI seed analysis on resting state data from 25 healthy normal adults. Seed maps from individual participants were used in a between-subjects ordinary least squares regression using FLAMEO, and the group-level statistical maps were subjected to multiple comparisons thresholding of Z > 3.1 and cluster correction of p < .05. This procedure created a thresholded map representing the mean functional connectivity of the disconnection mask across 25 healthy participants.

##### 2.3.2.2 Phase-scrambling analysis

In order to determine whether observed patterns of RSFC from the temporal pole seed were distinct from non-specific RSFC, we analyzed simulated phase-scrambled data. Our phase-scrambling procedure simulated data that shared the frequency composition of the observed data while randomizing phase information (Laumann et al., 2016). The disruption of phase information produced simulated timeseries that were not expected to produce the same RSFC patterns as the observed data despite closely resembling many attributes of the observed data. Our approach was implemented in custom Python software as follows. For each TD participant, we computed a fast Fourier transform on the mean RSFC time series from the disconnected mask (concatenated across all runs of data) to represent the timeseries in the frequency domain. Then — while preserving the observed amplitude in each frequency band — the phase component was randomly scrambled. Finally, an inverse fast Fourier transform was applied to the phase-scrambled frequency domain data in order to create a simulated timeseries in the time domain. This process was repeated to create 1000 simulated, phase-scrambled timeseries for each patient participant. These simulated time series had the same amplitude in each frequency band as the observed disconnected time-series, but were scrambled in phase from the real data, allowing us to map non-specific patterns of RSFC. Each of the phase-scrambled time series was correlated with every other voxel in the brain (as described above for the observed timeseries, ROI seed analysis) to produce 1000 phase-scrambled Fisher’s r-to-Z maps for each TD participant. These phase-scrambled maps were then tested against the observed disconnection correlation map using the SINGLETON option in AFNI’s 3DTTEST++ in order to compare the phase-scrambled distribution against the single observed TD RSFC map. The resulting t-statistic maps were transformed to Z-scores, and corrected for multiple comparisons using FSL’s CLUSTER command with a cluster defining threshold of Z > 3.1 and cluster correction of p < .05. This statistical map illustrated the extent to which the observed RSFC pattern of the time series data from the TD region differed from phase-randomized time series data having identical power spectra.

##### 2.3.2.3 ICA analysis

We also examined the RSFC data from our disconnection participants using a data-driven independent component analysis (ICA), as implemented in FSL’s Multivariate Exploratory Linear Optimized Decomposition into Independent Components (MELODIC) version 3.13 (Beckman & Smith, 2004). For each TD participant, the preprocessed, motion-scrubbed, concatenated 4D volume was used for the single-session ICA analysis. To guard against false positive results, we used an alternative hypothesis setting of p > 0.66 (which designates false-positive voxel assignment to be twice as bad as false-negative assignment relative to the default 0.5 even assignment) (FMRIB Analysis Group, 2013). We then used the disconnection mask for each participant to identify components with significant voxels within the disconnection mask. The fslcc tool from FSL was used to calculate the spatial correlation (thresholded at r > 0.2) between 10 well-replicated resting-state network maps identified and functionally characterized in Smith et al., 2009^1^ and TD participant components. We then identified TD participant components that were spatially correlated with the Smith et al. maps and had voxels in the disconnection mask for each TD participant.

##### 2.3.2.4 Comment on brain status and brain networks

The TD cases all had a history of pharmacoresistant temporal lobe epilepsy and long-term use of antiepileptic medications. Such factors could influence the organization and strength of functional brain networks. Prior work with epileptic populations has shown that epilepsy is associated with changes in brain networks relative to healthy comparisons (McGill et al., 2012; van Diessen et al., 2013; Waites, Briellmann, Saling, Abbott, & Jackson, 2006). Given our small sample size and the complex interactions of disease status with brain network structure, we did not have strong predictions regarding the pattern of any RSFC observed for TD cases beyond our main prediction that we would not observe RSFC between the disconnected temporal pole and other brain regions.

## 3. Results

### 3.1 Functional MRI, region of interest (ROI) seed analysis

We evaluated RSFC in TD cases using the disconnection mask that was made for each TD participant (Figure 1). Following from our hypothesis, we expected that for each of the five cases, we would not observe statistically significant RSFC with tissue outside the disconnected temporopolar tissue. Counter to our prediction, we observed significant RSFC between the disconnected temporal pole tissue and cortical areas in both hemispheres (Figure 2). This pattern of significant connectivity was observed at both stringent (Z > 3.1) and more liberal cluster forming thresholds (Z > 2.33). The extent of RSFC in each case varied, but we did observe some common patterns. Two cases, 3360L and 3753R, showed RSFC between the disconnected pole and bilateral medial frontal regions, similar to the pattern of RSFC seen in healthy comparison data. In addition, two cases, 3749L and 3786L, showed RSFC with non-resected bilateral hippocampal areas (though only at the more liberal cluster threshold). 3749L showed connectivity in the anterior hippocampus, and 3786L showed connectivity in the posterior hippocampal areas. 3749L, as well as 3794R, also showed RSFC between the disconnected tissue and regions in the contralateral, non-resected temporal pole. Some cases showed RSFC between the disconnected temporopolar tissue and brain regions that were not present in the corresponding healthy RSFC map. For example, 3786L showed connectivity with parts of the occipital pole that were not observed in the healthy RSFC map. 3753R and 3794R showed connectivity with portions of the posterior cingulate cortex that do not appear in the map of the disconnected tissue seeded in healthy comparison data. The coordinate reports for peak clusters in the healthy comparison and TD cases are reported in Tables S2, S3 and S4.

**Figure 2:**
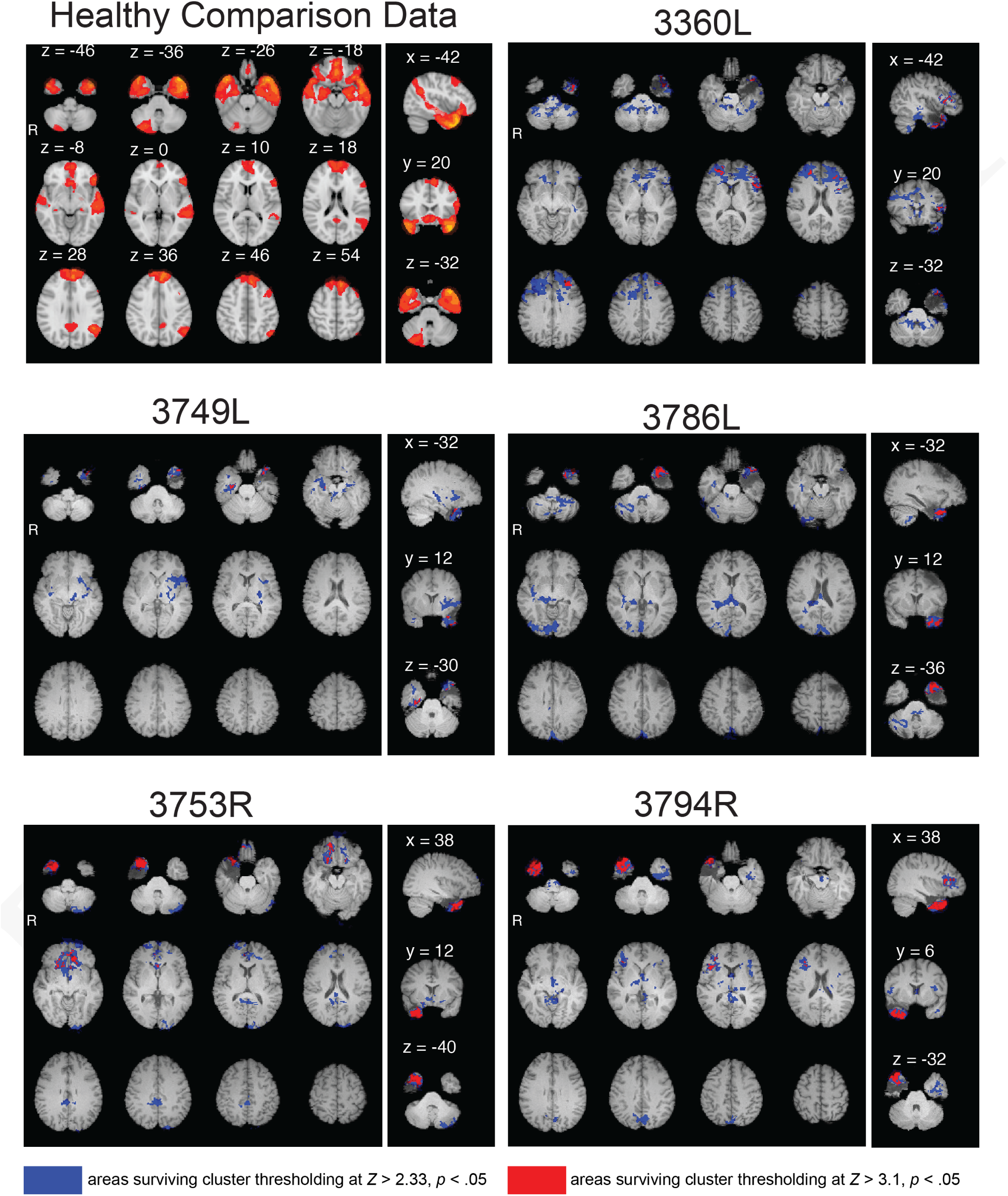
Functional connectivity maps showing significant connectivity from each participant’s disconnected tissue mask. Upper left panel shows pattern of connectivity from 3360’s disconnection mask seed in 25 healthy adults. Regions in blue represents cluster thresholding at Z > 2.33, p < .05, and regions in red indicate those areas surviving cluster thresholding at Z > 3.1, p < .05. Coordinate labels represent MNI152 2mm atlas space slices. For peak cluster coordinates, see Tables S2, S3, and S4.

The observation of significant, bilateral RSFC in each of the five TD cases went against our prediction that there would be no RSFC outside of the disconnected temporal pole.

### 3.2 Functional MRI, phase-scrambled analysis

To evaluate whether the observed patterns of RSFC could be explained solely by chance, an additional analysis was conducted in which we simulated many timeseries that had the same power spectrum as the observed TD timeseries but scrambled phase. As illustrated in Figure 3A, substantial portions of the brain exhibited RSFC with the observed TD timeseries versus the simulated phase-scrambled data. When comparing these results against the cluster-thresholded fixed-effects seed maps (Figure 3B), we observed that the pattern of specific TD connectivity was highly overlapping with the phase-scrambled analysis. The convergence of these two unique analyses suggests that the observed pattern of RSFC is not a simple statistical artifact. The coordinate reports for peak clusters in the TD vs phase-scrambled analysis are reported in Table S5. Findings from this analysis also went against our hypothesis that the disconnected temporal pole would not show significant RSFC with other parts of the brain.

**Figure 3:**
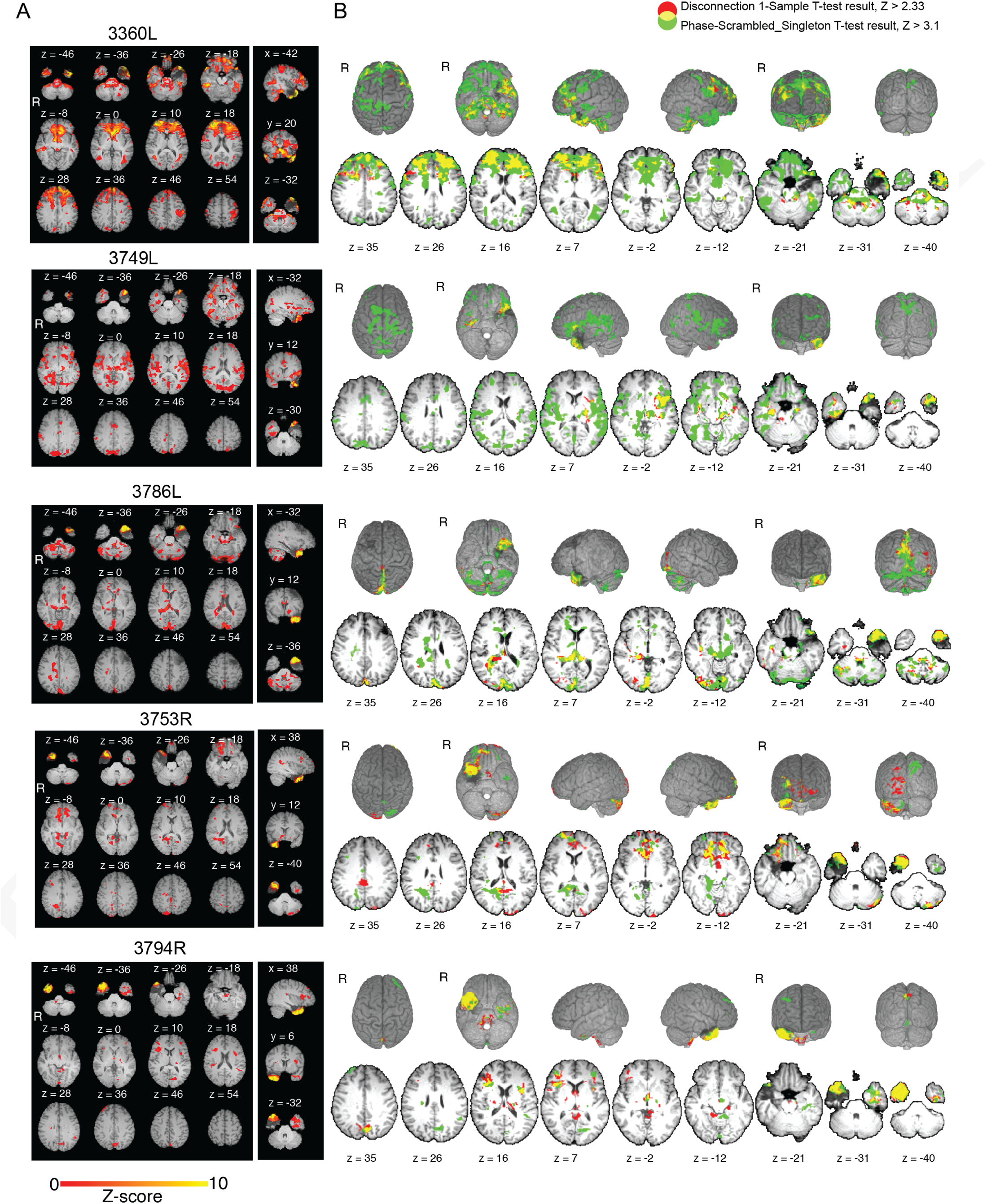
Functional connectivity maps showing significant connectivity for each participant’s disconnected tissue mask compared to 1000 simulated phase-scrambled maps, cluster thresholded at Z > 3.1, p < .05 (A). Connectivity overlap of fixed-effect maps (shown in Figure 2C) (in red) and phase-scrambled contrast (in green) are shown in (B). Overlaps of the two maps appear in yellow. For peak cluster coordinates, see Table S5.

### 3.3 Functional MRI, ICA analysis

We also investigated the RSFC of the TD cases using a data-driven ICA analysis. In each TD participant, we identified several ICA-derived components that contained significant voxels within the disconnected tissue. Within these components, we examined the degree to which the spatial pattern of these components matched 10 well-replicated networks identified in healthy participants (Smith et al., 2009), and we observed several components showing good correspondence (Figure 4). Of note, 3749L and 3360L had components with high spatial similarity with known medial (3749L component 2, r = 0.774) and lateral (3360L component 4, r = 0.709) visual networks, respectively, while 3786L and 3753R had components that were similar to the frontopari-etal network (3786L component 6, r = 0.511; 3753R component 5, r = 0.453). Finally, 3794R had at least one component showing good correspondence with the default mode network (component 31, r = 0.478). A full list of components with significant connectivity in disconnected pole and spatial correlations with Smith et al. (2009) components is provided in Table S6. In summary, we observed data-driven components including connectivity with the disconnected tissue that were spatially correlated with separately identified resting-state networks in healthy participants, providing further evidence that the connectivity observed with the disconnected temporal pole is not simply artifact.

**Figure 4:**
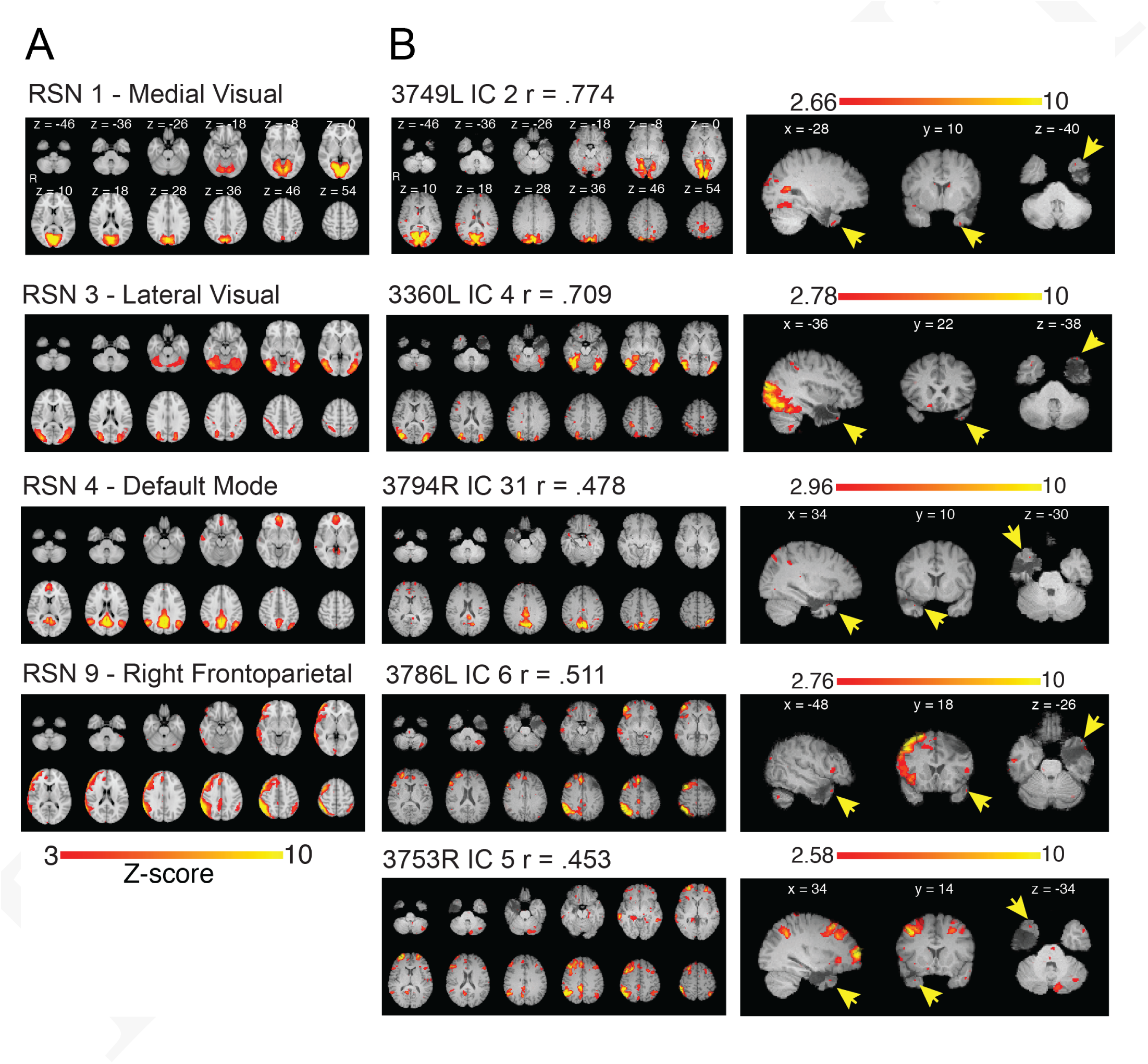
Correspondence between well-replicated resting-state network maps identified by Smith et al. 2009 (A), and independent component maps from disconnection participants (B). The correlation reported is the spatial correlation between the Smith et al. 2009 map and the disconnected participant component. Significant voxels in the disconnected tissue are highlighted by yellow arrows. Coordinate labels represent MNI152 2mm atlas space slices. For a full list of correlated components, see Table S6.

## 4. Discussion

Our findings have significant implications for studies using functional connectivity measures because — for the first time — we show that a brain region confirmed to have no structural neural connections can show reliable functional connectivity with remote brain regions. These results challenge a key premise in the field: that functional connectivity reflects mono-or polysynaptic communication along structural pathways. In addition, these results strongly suggest an underappreciated role of other physiological contributors to large-scale variations in BOLD signal across the brain that are unaccounted for in current analysis standards. Our current investigation was not designed to arbitrate which mechanism or mechanisms might have contributed to our findings, but we believe that our observations underline the importance of ongoing basic science addressing the origins of functional connectivity as studied with fMRI.

Our findings of RSFC in the human brain between regions that have been completely surgically disconnected are novel, although there have been prior reports of preserved functional connectivity in humans despite significant anatomical disruption (Lu et al., 2011; Tyszka et al., 2011; Uddin et al., 2008). Patients who lack a corpus callosum due to surgical intervention or abnormal development have been shown to retain essentially normal interhemispheric functional connectivity (Tyszka et al., 2011; Uddin et al., 2008), but it has been noted that many structural pathways still join the two hemispheres in these patients (e.g., anterior and posterior commissures, midbrain, brainstem). Although these non-resected connections are sparse or indirect in comparison to the corpus callosum, the possibility remains that they could continue to facilitate interhemispheric connectivity (O’Reilly et al., 2013). Beyond intentional resection, structural disconnection can also occur because of naturally-occurring etiologies such as stroke. Here too, structural disconnection has been shown to exert a surprisingly moderate influence on normal patterns of functional connectivity (Lu et al., 2011). While these earlier findings implied that non-obvious mechanisms might contribute to functional connectivity, in no previous study was physical disconnection of a brain region microscopically verified in vivo during a resection procedure.

In conjunction with prior work, our findings cast doubt on the premise that structural connectivity is necessary for functional connectivity as observed with fMRI. Our report has the advantage of a strict baseline condition — in vivo RSFC measures from surgically-disconnected brain tissue — but other authors have drawn similar conclusions from alternative evidence. Using MRI measures of healthy structural and functional connectivity, Honey et al. (2009) observed that structural and functional connectivity were correlated (Pearson’s r=0.46-0.70) but the relationship left significant additional variance unexplained. Sources of non-neuronally originated functional connectivity in rs-fMRI include physiological signals such as cardiac and respiratory rhythms (Weisskoff et al., 1993), head motion (Power et al., 2012; Satterthwaite et al., 2012; Koene R. A. Van Dijk et al., 2012), low-frequency drift (Shmueli et al., 2007) and others. Critically, we applied standard remedies for each of these noise variables during data (pre)processing to reduce contributions of spurious signals to our observed RSFC (see Materials and Methods). Thus, while there is strong evidence suggesting that structural connectivity contributes significantly to functional connectivity, current approaches are insufficient to control for noise variables and/or other mechanisms must also contribute to functional connectivity in the BOLD signal as observed with fMRI.

### 4.1. Putative mechanisms for functional connectivity of disconnected temporal pole

We offer three possible mechanisms that could account for our findings, each of which draws on literature describing factors regulating cerebral blood flow (CBF). First, there is empirical evidence that coordinated vasodilation can span surprisingly long distances (Osol & Halpern, 1988; Porret, Stergiopulos, Hayoz, Brunner, & Meister, 1995). Temporal patterns in vascular resistance variation (i.e., constriction and relaxation of arterial walls) are complex and have been described as chaotic, non-linear, and not strongly related to systemic factors such as blood pressure or heart rate (Stergiopu-los, Porret, De Brouwer, & Meister, 1998). In spite of this complexity, very strong correlations have been observed in arterial diameter oscillations between discrete arteries in one or both upper limbs of human participants (Porret et al., 1995). Critically, these patterns did not exhibit phase progression that would imply a slowly spreading signal, but were instead phase-locked in a manner suggestive of widespread vascular coordination. Explanatory mechanisms that could support this coordination are currently being investigated but remain unknown. For example, it has been reported that local capillary beds may provide feedback regarding perfusion in a rapid and ongoing stream by sending signals through gap junctions in the endothelium lining each vessel (Bagher & Segal, 2011; Haddock & Hill, 2002). We note that these vascular changes are distinct from physiological factors that have been frequently discussed in the fMRI literature including cardiac and respiratory signals (Shmueli et al., 2007; Weisskoff et al., 1993). Here, any influence of these signals should have been minimized by our data processing approach which incorporated recommended prophylactic steps (e.g., bandpass filtering, global signal regression, white matter signal regression) (K. R. A. Van Dijk et al., 2010; Power, Schlaggar, & Petersen, 2014). However, subtle vascular influences on the fMRI measures may be less affected by these remedies.

While the vasculature of the entire body displays a surprisingly robust response to local changes in perfusion, a second possible mechanism for our findings is the strong influence of direct stimulation by neural sources on the vasculature of the brain (Hamel, 2006). In particular, nuclei of the brainstem and basal forebrain have been shown in animal models to exert widespread influence on CBF (Bekar, Wei, & Nedergaard, 2012; Hotta, Uchida, Kagitani, & Maruyama, 2011; Toussay, Basu, Lacoste, & Hamel, 2013; Vaucher & Hamel, 1995). Noradrenergic signals originating from the locus coeruleus enhance functional hyperemia and may serve to optimize neurovascular coupling (Bekar et al., 2012). Empirically, stimulation of the locus coeruleus unilaterally produces ipsilateral enhancement of CBF throughout substantial portions of the forebrain (Toussay et al., 2013). Meanwhile, portions of the basal forebrain also exercise some control over CBF, whether through cholinergic pathways driven by the nucleus basalis of Meynert (Hotta et al., 2011) or through alternative routes driven by the non-cholinergic neurons in the substantia innominata (Vaucher & Hamel, 1995). These diverse nuclei have each been shown to influence CBF in isolation, but in the context of a normally functioning brain they may cooperate to maintain optimal conditions in response to regional differences in brain activity. Critically, the collective influence of these nuclei on CBF appears to reach nearly the entire brain, and these nuclei may therefore provide some degree of central coordination and potentially originate certain synchronized changes in CBF. Thus, it is possible that the influence of these (and potentially other) brain regions that globally affect CBF could be providing an exogenous signal to preserved local vasculature near TP which is unaffected by focal injury (including structural disconnection) and that is sufficient to support RSFC.

Neural signals from within the central nervous system affect CBF, but preserved autonomic innervation of the pial vasculature provide a third potential explanatory mechanism. Studies of the peripheral nervous system have shown that autonomic ganglia associated with certain cranial nerves can affect CBF of the brain (Boysen, Dragon, & Talman, 2009; Taktakishvili, Lin, Vanderheyden, Nashelsky, & Talman, 2010; Talman, Corr, Nitschke Dragon, & Wang, 2007; Talman & Nitschke Dragon, 2007). Post-ganglionic parasympathetic fibers from the pterygopalatine ganglion innervate nearby cerebral blood vessels, and these fibers have been reported to exert a tonic influence on CBF in the forebrain (Boysen et al., 2009; Talman et al., 2007). Removal of this innervation immediately reduced CBF by approximately 20%, indicating a substantial and unexpected role of parasympathetic ganglia in maintaining CBF in the brain. The RSFC that we observed could therefore reflect the activity of parasympathetic mechanisms, but notably these parasympathetic ganglia are by no means isolated from the CNS. The pre-ganglionic parasympathetic neurons are located in nuclei of the pons and medulla, and these nuclei are in turn influenced by many ascending and descending pathways. In addition, sympathetic mechanisms can also affect CBF as demonstrated by the effects of resecting or stimulating the superior cervical ganglion and thereby altering its innervating influence on the internal carotid artery (Gross, Heistad, Strait, Marcus, & Brody, 1979; Sato et al., 1990). In combination, these sympathetic and parasympathetic mechanisms provide further means — albeit indirect — by which the CNS can influence CBF which could potentially create unexpected RSFC of structurally disconnected brain regions.

The presence of confounding variables in the collection or analysis of functional neuroimaging data is another possibility, but we attempted to mitigate these by rigorously applying published artifact reduction techniques that are widely employed in the field. Likewise, residual brain neural connections in the gray or white matter of the cerebral cortex could drive the results we found, but no such connections were evident at surgical or postsurgical examinations. We argue that (1) our neuroimaging data reflect non-spurious BOLD-related functional connectivity associated with the temporal poles and (2) the temporal pole of each participant was fully disconnected from cortical gray and white matter, and we have identified several explanatory mechanisms that could potentially account for the findings. These may include large-scale coordination of cerebral blood flow via mechanisms controlling vascular resistance, such as central control of autonomic vasomotor activity or direct innervation by deep brain nuclei. Our current study cannot arbitrate among these or other possible explanations, but further research in human participants and animal models may provide useful insights.

### 4.2 Limitations

Our study had some limitations. The number of TD participants was relatively small, due to the relative rarity of this type of resection in our Registry. Concerns about statistical power are especially pertinent with small sample sizes, but we note that our findings were described at the individual level in all cases in order to address this concern and that our key findings are positive rather than null. Notably, positive findings were not related to the time since resection as illustrated by case 3360L whose data still showed significant functional connectivity of disconnected temporal pole a decade after surgery. Also, as discussed elsewhere, fMRI signal from the temporal region is prone to susceptibility artifacts (Devlin et al., 2000; Power et al., 2011; Weiskopf, Hutton, Josephs, & Deichmann, 2006; Yeo et al., 2011), but this same constraint would apply to a substantial majority of studies examining temporal lobe functional connectivity based on whole-brain functional images (e.g., Andrews-Hanna, Reidler, Sepulcre, Poulin, & Buckner, 2010). In the event that unknown or poorly characterized noise variables (whether intrinsic or extrinsic) drove our current findings, other investigations that employ typical methods for fMRI data collection, processing, and analysis may be susceptible to the influence of the same variables. Also similar to many neuroimaging studies, we did not have access to physiological data from the fMRI scanning sessions that might have reduced any remaining influence of physiological variables on the resting-state BOLD signal. Another limitation was that our study relied solely on structural and functional MRI data, and other neuroimaging techniques might prove informative. We infer that the tissue of the disconnected temporal pole persists because of intact vascular supply, but PET-based perfusion data might substantiate this inference. Similarly, we assume that the functional status of the same tissue is abnormal and not related to that of other brain regions, but intracranial recordings could potentially establish this empirically. Notably, the temporal focus of each TD case’s epilepsy may have led to atypical organization of functional connections with other brain regions even prior to surgery (cf. van Diessen, Diederen, Braun, Jansen, & Stam, 2013 for a discussion of brain network changes in chronic epilepsy), but we suggest that abnormal functional connections would still not be expected to persist after surgery. To summarize, our study has very important, albeit initial, findings from a line of investigation that our laboratory and others will continue to pursue.

### 4.3 Conclusions

The study of functional connectivity has provided converging evidence regarding the neural correlates of cognitive processes, disease processes, and functional networks (Buckner et al., 2013; Power et al., 2011; Sabuncu et al., 2011; Yeo et al., 2011; Zhang & Raichle, 2010). Previous focused critiques of typical fMRI-based statistical methods (Bennett, Wolford, & Miller, 2009; Eklund, Nichols, & Knutsson, 2016), potential sources of noise (Birn, Diamond, Smith, & Bandettini, 2006; Devlin et al., 2000; Leopold & Maier, 2012; Power et al., 2014; Shmueli et al., 2007; Sirotin & Das, 2009; Weiskopf et al., 2006; Wise, Ide, Poulin, & Tracey, 2004), or interpretation of results (Poldrack, 2006; Vul, Harris, Winkielman, & Pashler, 2009) have provided important cautions about the importance of sound methodology. In the same vein, we hope that our findings will draw attention to underappreciated mechanisms influencing functional connectivity analysis.

We observed significant whole-brain resting-state functional connectivity in the fMRI BOLD signal obtained from the temporal pole despite the surgical disconnection of that brain region from the rest of the brain. This unexpected RSFC did not appear to be due to residual anatomical connections, and our findings challenge an important premise underlying conventional interpretations of functional connectivity analysis. Despite the cryptogenic origin of the observed RSFC, we believe that a variety of potential mechanisms may contribute to the coordinated activation of physically disconnected brain regions including several sources of direct or indirect influence on CBF. Determining which mechanism or mechanisms directly support this outcome is for future study, and methods beyond fMRI may be informative in this regard. Until such investigations elucidate this topic, our results provide a heretofore non-observed baseline measurement for an increasingly popular neuroimaging technique. While we do not believe that the observed patterns of functional-without-structural connectivity pose a direct challenge to the application of functional connectivity methods, we do believe that these findings provide a clear illustration of the long acknowledged, but sometimes underappreciated, interplay of neural and vascular signals underlying BOLD-based fMRI analysis methods. Our results reinforce the importance of cautious interpretation of findings from RSFC investigations, particularly in the absence of convergent evidence from task-based fMRI, behavior, and animal studies.

## 5. End Matter

### 5.1 License

© 2017 by the authors. All rights reserved.

## 5.2 Acknowledgements

We would like to thank the participants and their families for their contributions to this project. Additionally, we wish to thank the following individuals for helpful commentary on this project: William Talman and Patrick Drew.

## 5.5 Author Contributions

All authors designed the research; DEW, MJS, JB, TJA, & AJ performed the research and conducted the analyses; all authors wrote the paper. All authors granted permission for this preprint to be posted to BIORXIV.

## Footnotes

^1^http://www.fmrib.ox.ac.uk/datasets/brainmap+rsns/

